# AbRefine: Framework-Conditioned Refinement for Structure-Guided Antibody CDR Design

**DOI:** 10.64898/2026.09.27.754743

**Authors:** Qi Liu, Xinyi Xu, Qian Cheng, Chen Wu, Lu Meng, Defu Yang, Xinbo Gao, Xueli Chen

## Abstract

Structure-guided antibody CDR design constrains residue selection using backbone geometry, but structural evidence alone may leave multiple amino acids plausible. We investigate whether the fixed antibody framework provides complementary sequence information for resolving this ambiguity. We introduce AbRefine, an information-aware fusion framework that combines backbone-conditioned structural predictions with framework-conditioned sequence distributions derived from a pretrained antibody language model with all designed CDRs masked. Its Information-Aware Fusion (IAF) module learns residue-specific fusion weights from the uncertainty, disagreement, and information content of the two predictive distributions, while keeping both pretrained experts frozen. Controlled interventions show that the signal is target specific: matched frameworks outperform shuffled frameworks by 7.37 percentage points, and the largest gain occurs at positions with high structural uncertainty and informative framework evidence (+5.77 points). Compared with static fusion, IAF further improves AbMPNN recovery by +1.53 points (95% CI +0.93 to +2.14) and significantly reduces native negative log-likelihood for both AbMPNN and AntiFold, without detectable degradation in fold-back structural compatibility. These results show that framework context provides information beyond backbone geometry and that learning its residue-specific utility enables effective adaptive fusion for antibody CDR design.

## 1. Introduction

Antibody optimization often begins with an established molecule whose scaffold and binding-site geometry are already known or desired. In this setting, redesigning complementarity-determining regions (CDRs) is not an unconstrained sequence-generation problem: candidate residues should remain compatible with the prescribed backbone and the surrounding framework. This setting arises when exploring sequence alternatives around an existing lead, adapting CDRs to a selected framework, or optimizing an antibody while seeking to retain an established binding geometry. Importantly, framework–CDR interactions can influence loop conformations and antigen recognition, making the framework an active part of the design context rather than a passive scaffold [2, 9].

Structure-guided sequence design provides a natural way to enforce geometric compatibility. Fixed-backbone models infer aminoacid preferences from a prescribed structure and have been successfully extended from general proteins to antibody CDRs [5, 11]. Yet a backbone does not necessarily specify a unique sequence: multiple amino acids may remain compatible with similar local environments. The resulting *residual sequence ambiguity* is particularly relevant in CDRs, where residue preferences depend not only on geometry but also on the antibody sequence background. Thus, once structural conditioning has narrowed the set of plausible residues, an important question remains: what additional information can distinguish among the alternatives left unresolved by geometry?

We investigate the fixed antibody framework as such an information source. Antibody language models learn contextual dependencies from large repertoire collections and can estimate residue preferences from sequence context alone [15, 16]. Recent work has further shown that language-model predictions can complement structure-based antibody design [6]. However, this raises two unresolved questions: whether the useful sequence signal originates specifically from the *target framework*, and whether that signal should influence every design position equally. By masking all designed CDR residues, the framework can be isolated as an observable sequence context, providing a framework-conditioned distribution that is distinct from the backbone-conditioned structural distribution.

The second question leads naturally to an information-fusion problem. Additional information is useful only when it contributes predictive evidence beyond what is already available. Information-theoretic studies of multimodal fusion characterize this utility in terms of uncertainty, redundancy, and disagreement between information sources [4], while knowledge-enhanced learning has shown that indiscriminate incorporation of auxiliary information can be ineffective when that information is unreliable [19]. Related work on knowledge amalgamation further demonstrates the value of selectively integrating heterogeneous pretrained models [10], and recent biological applications increasingly use adaptive mechanisms to combine heterogeneous molecular information [13]. These observations motivate a residue-specific rather than globally fixed treatment of framework information.

To address this problem, we introduce **AbRefine**, an information-aware fusion framework for structure-guided antibody CDR design. AbRefine combines a backbone-conditioned structural expert with a framework-conditioned antibody language model while keeping both pretrained models frozen. Its *Information-Aware Fusion* (IAF) module characterizes each residue through the information state of the two predictive distributions—including their uncertainty, disagreement, and framework-derived information content—and learns a residue-specific fusion weight. Framework evidence can therefore contribute strongly when it provides useful complementary information and remain limited when the structural prediction is already sufficient.

Our experiments provide evidence for both components of this formulation. Controlled framework interventions show that target-matched frameworks contain CDR-relevant information that is lost after framework shuffling and cannot be reproduced by generic CDR residue priors. The utility of this information is strongly position dependent, with the largest gains occurring where structural predictions are uncertain and framework evidence is informative. IAF translates this relationship into adaptive fusion, providing additional gains over static fusion when substantial residual ambiguity remains and improving native likelihood across structural experts without detectable degradation of fold-back structural compatibility.

Our contributions are threefold:

- We identify the fixed antibody framework as a source of target-specific information that complements backbone geometry in CDR residue selection, particularly where structural predictions remain ambiguous.
- We introduce **AbRefine** with Information-Aware Fusion (IAF), which learns residue-specific fusion weights from the information state of structure- and framework-conditioned distributions, enabling adaptive integration of two frozen pretrained experts.
- Through controlled framework interventions, cross-expert evaluation, and information-utility analysis, we characterize when framework information improves CDR design and show that adaptive fusion provides additional benefit when the structural expert leaves substantial residual ambiguity.

## 2 Related Work

### 2.1 Structure-guided antibody design

Fixed-backbone sequence design seeks amino-acid sequences compatible with a prescribed protein geometry. ProteinMPNN introduced an efficient message-passing architecture for this setting, while ESM-IF1 learned structure-conditioned sequence representations from large collections of predicted structures [5, 12]. Antibody-specific models subsequently adapted fixed-backbone design to immunoglobulin structures. AbMPNN specializes ProteinMPNN for antibody sequence design, whereas AntiFold fine-tunes ESM-IF1 on antibody structures and improves sequence recovery in hypervariable CDRs [7, 11].

A related line of work jointly models antibody sequence and geometry rather than holding the target CDR backbone fixed. DiffAb formulates antigen-conditioned CDR design as diffusion over sequence and structure [14]. FlowDesign replaces diffusion with flow matching and improves the treatment of sequence and structural priors [18], while GeoGAD incorporates multi-scale geometric representations for CDR design and affinity optimization [17]. RFdiffusion-based antibody design further conditions on an antibody framework while generating CDR conformations and antibody– target binding poses [3]. These methods span fixed-backbone redesign and increasingly constrained generation of CDR sequence, conformation, and binding geometry.

### 2.2 Antibody language models and sequence guidance

Antibody language models learn sequence regularities from large repertoire collections independently of three-dimensional structure. AntiBERTy learns contextual representations associated with antibody repertoire organization and affinity maturation [16], while AbLang uses masked language modeling to reconstruct antibody sequences from surrounding context [15]. These models have established pretrained sequence representations as a useful source of antibody-specific information.

Sequence and structural models have also been combined for antibody design. Del Alamo et al. ensemble ProteinMPNN with AbLang and report improved CDR design, including substantially more experimentally recovered HER2-binding CDRH3 variants than with ProteinMPNN alone [6]. Such results establish the utility of combining antibody language-model predictions with backbone-conditioned design. Existing combinations, however, primarily treat sequence-model output as an additional design prior or ensemble component rather than explicitly modeling how the utility of that information varies across individual design positions.

### 2.3 Adaptive information fusion

Information fusion aims to combine heterogeneous sources whose predictive value, redundancy, and reliability may vary across observations. Information-theoretic analyses characterize multimodal fusion in terms of the additional information contributed by each source and the effects of uncertainty, redundancy, and conflicting evidence [4]. In knowledge-enhanced biomedical learning, auxiliary domain information has likewise been shown to be beneficial only when it is sufficiently reliable, motivating selective rather than indiscriminate incorporation [19].

Related work has developed learned mechanisms for combining heterogeneous models and representations. Knowledge-amalgamation approaches integrate information from multiple pretrained teachers without requiring access to their original training data [10], while multimodal biological models combine heterogeneous molecular and protein representations through learned fusion mechanisms [13]. Most such approaches perform fusion at the feature or representation level, where task-specific networks learn interactions among latent embeddings.

AbRefine considers a different fusion setting. Its structural and sequence experts are independently pretrained and already produce normalized distributions over the same amino-acid output space. Rather than learning a joint representation, Information-Aware Fusion operates directly on these predictive distributions and estimates a residue-specific fusion weight from their information state. This positions AbRefine as an adaptive distribution-level fusion method for combining structural and framework-conditioned sequence evidence in antibody CDR design.

## 3 Method

### 3.1 Problem formulation and conditional experts

We consider CDR sequence design under a prescribed antibody backbone and a fixed framework sequence. Let *X* denote the paired heavy–light backbone, *F* the observed framework residues, and *C* = (*c*_1_, …, *c*_*L*_) the CDR residues to be designed. The objective is to model

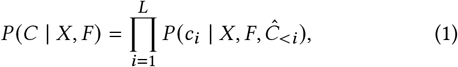

using only information available at design time. Native CDR identities are never provided as model inputs.

AbRefine represents each design position using two predictive distributions over the amino-acid alphabet *A* . The structural expert provides

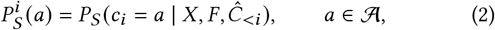

where *Ĉ*_*<i*_ denotes the CDR prefix available to the structural model at position *i*. AntiFold is used as the primary structural expert, and additional experiments instantiate the same formulation with AbMPNN. All structural-model parameters remain frozen.

The second expert extracts sequence information from the observed framework without exposing the CDR identities being designed. For an antibody chain with sequence *S* = (*s*_1_, …, *s*_*n*_ ), we construct

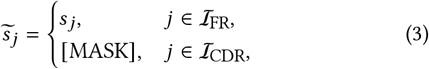

where *I*_FR_ and *I*_CDR_ denote framework and CDR positions, respectively. AntiBERTy is applied separately to each chain: H1, H2, and H3 are masked simultaneously on the heavy chain, and L1, L2, and L3 are masked simultaneously on the light chain. It then provides

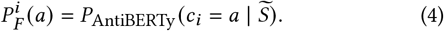

Thus, 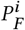 depends on the retained same-chain framework and the pretrained language model, but not on native CDR identities, backbone coordinates, or generated CDR residues. The AntiBERTy parameters are also kept frozen.

The two experts provide heterogeneous evidence over the same residue variable. 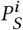 represents compatibility with the prescribed structural context, whereas 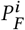 represents sequence preferences conditioned on the observed framework. AbRefine learns how strongly the latter should modify the former at each design position.

### 3.2 Information-Aware Fusion

For each CDR residue, we summarize the state of the two predictive distributions using four quantities. Their uncertainties are measured by entropy,

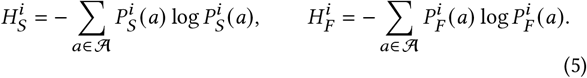

Cross-expert disagreement is measured as

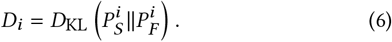

We additionally compare the framework-conditioned distribution with a generic development-derived CDR prior. For each of H1, H2, H3, L1, L2, and L3, CDR positions are mapped to ten bins according to normalized loop position. Amino-acid frequencies within each CDR–position bin are estimated with a pseudocount of 0.5 per amino acid, yielding 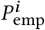. During development, the empirical prior for each antibody is constructed after excluding that complete antibody. For evaluation targets, *P*_emp_ is constructed once from all 62 development antibodies and then frozen; no test sequence contributes to the prior.

We define

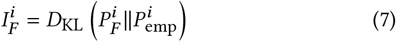

as an informativeness proxy. 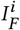 measures how strongly the framework-conditioned language-model distribution departs from a generic development-derived CDR distribution; it is not interpreted as a direct estimate of causal target-specific framework information. Target specificity is evaluated separately through matched-versus-shuffled framework interventions.

The resulting information-state vector is

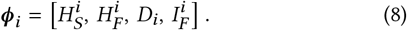

Because these quantities have different numerical scales, they are standardized using statistics computed exclusively from the development antibodies:

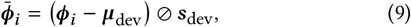

where *µ*_dev_ and ***s***_dev_ are the development-set feature means and standard deviations, and ⊘ denotes element-wise division. These statistics are frozen before evaluation.

IAF is parameterized as a residue-specific correction around a globally calibrated fusion point rather than learning fusion weights from scratch. Let *g*_0_ = 0.59 denote the fixed product-of-experts coefficient selected on the independent development set. For ensemble member *k*,

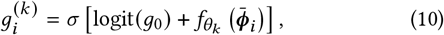

where 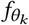 is a 4 8 1 multilayer perceptron with a SiLU hidden activation and 49 trainable parameters. The output layer is initialized to zero, so the initial model satisfies 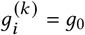 for every residue. IAF therefore learns residue-specific deviations from the development-selected static fusion rule.

Each ensemble member combines the two predictive distributions as

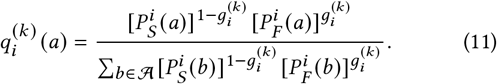

Fusion is implemented in log-probability space for numerical stability. We train three IAF models with seeds 41, 42, and 43. The reported refined distribution averages their *probability outputs*,

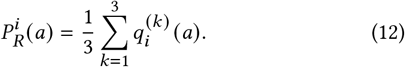

We average the three distributions rather than the gate values because the nonlinear fusion in Eq. (11) makes these operations non-equivalent.

### 3.3 Properties of information-aware fusion

The adaptive product-of-experts formulation admits a simple interpretation of why residue-specific fusion is useful. For clarity, we consider one ensemble member and omit the residue and seed indices. Let

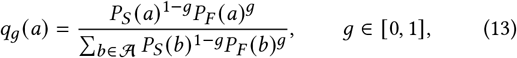

and define the expert log-ratio

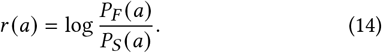

Equation (13) can then be rewritten as

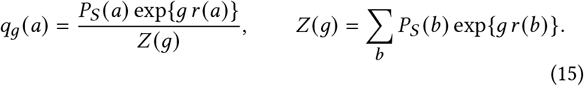

Thus, for fixed expert distributions, adaptive fusion forms a one-parameter exponential family in which *g* controls how strongly the framework expert reweights the structural distribution.

*Residue-specific fusion has a well-defined local objective*. For the native residue *y*, let

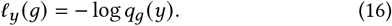

Differentiating with respect to *g* gives

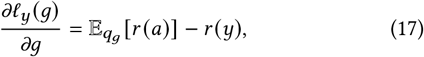

and

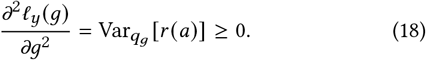

Hence, the native-residue NLL is convex in the fusion coefficient for fixed *P*_*S*_ and *P*_*F*_ . In particular, increasing framework influence locally improves the native likelihood whenever

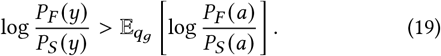

Conversely, when the framework expert provides relatively weaker support for the native residue, decreasing its contribution is locally preferable. This shows why a single global coefficient need not be optimal for every residue: different expert pairs can induce different residue-level fusion optima.

*Expert disagreement determines sensitivity to fusion*. Equation (18) also characterizes how strongly the fused distribution can respond to changes in *g*. If the two experts agree exactly, *P*_*S*_ = *P*_*F*_, then *r* (*a*) = 0 for all *a* and

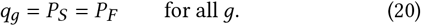

More generally, when the experts are similar, the variance of *r* (*a*) is small and changing the fusion coefficient has limited effect. Substantial disagreement creates greater scope for adaptive reweighting. Structural uncertainty alone is therefore insufficient to determine fusion utility: a structural expert can be uncertain while the framework expert provides redundant or uninformative evidence. This motivates jointly characterizing uncertainty and cross-expert disagreement in IAF.

*IAF performs regularized local adaptation around a global solution*. IAF does not learn the fusion coefficient from an arbitrary initialization. Its gate is parameterized as

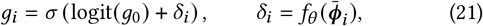

where *g*_0_ = 0.59 is selected on the development set. A first-order expansion around δ_*i*_ = 0 yields

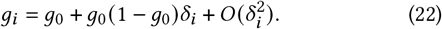

Thus, near initialization, IAF learns a smooth residue-specific correction to the globally calibrated operating point rather than relearning fusion from scratch. The regularization term

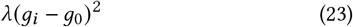

further penalizes unsupported departures from this anchor.

Together, these properties motivate the information state used by IAF. Structural entropy *H*_*S*_ describes the need for additional evidence, framework entropy *H*_*F*_ describes the concentration of the auxiliary prediction, and *D*_KL_ (*P*_*S*_ ∥ *P*_*F*_ )measures non-redundancy between the two experts. The fourth feature,

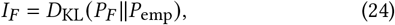

measures the departure of the framework-conditioned distribution from a generic development-derived CDR prior and is used only as an informativeness proxy. It does not by itself establish target-specific framework information, which is tested separately through matched-versus-shuffled framework interventions.

The resulting interpretation is that IAF approximates residue-specific fusion utility from quantities available at inference time: uncertainty indicates where refinement may be needed, disagreement determines whether fusion can materially alter the prediction, and framework-distribution specificity helps characterize whether the auxiliary evidence departs from generic CDR preferences. The anchored gate then converts this information state into a regularized local correction of the global fusion rule.

### 3.4 Training and decoding

IAF is trained only on the development antibodies; neither pretrained expert is updated. Let the development collection contain *M* antibodies and let *I*_*t*_ denote the evaluated CDR residues of antibody *t* . To prevent antibodies with more CDR residues from receiving greater training weight, the likelihood term is computed within each antibody and then macro-averaged across targets:

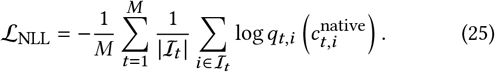

We regularize the learned gates toward the static fusion anchor,

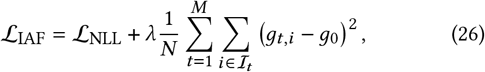

where *N* = Σ_*t*_ |*I*_*t*_|.Five-fold antibody-grouped development validation selects λ = 0.01; complete antibodies remain within a single fold throughout model selection. After hyperparameter selection, three models with seeds 41–43 are trained on the full development collection and frozen for all subsequent evaluations. The anchor *g*_0_ = 0.59, empirical prior, feature-standardization statistics, and all IAF parameters are determined without using evaluation targets.

We consider two decoding protocols. In *open-loop* evaluation, the structural distributions 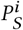 are evaluated along the original frozen structural-model trajectory, and AbRefine refines each residue distribution without writing its prediction back into subsequent structural contexts. This setting isolates the immediate effect of information-aware fusion from changes in the autoregressive trajectory.

In *closed-loop* decoding, the refined distribution is used to select

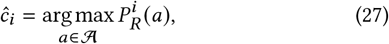

and *ĉ*_*i*_ is written back before the structural expert predicts the next CDR position. The resulting autoregressive model can be written as

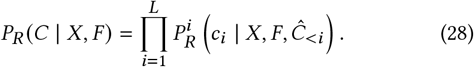

The framework-conditioned distributions are computed once from the all-CDR-masked heavy or light chain and remain cached and unchanged in both decoding modes. Consequently, generated CDR residues affect subsequent predictions only through the structural expert, while the sequence expert continues to represent the original framework context.

## 4 Experimental Setup

### 4.1 Datasets and evaluation cohorts

All model development is restricted to 62 SAbDab antibodies comprising 3,077 CDR residues. These targets are obtained from an initial collection of 75 experimental antibody–antigen complexes after removing 12 antigen-family overlaps and one exact-PDB overlap with RAbD-Epi. Antibodies are grouped using 90%-identity heavy-chain and antigen families before assignment. The development collection is used for feature normalization, empirical-prior construction, hyperparameter selection, and IAF training; no evaluation target contributes to these quantities.

Our primary evaluation uses the public SAbDab test partition released with AbMPNN and subsequently used by AntiFold. Of 405 deduplicated test Fabs, 362 have reconstructable paired heavy– light records. Removing three exact-PDB overlaps with the development collection (5w3l:EG, 5w3m:EG, and 5w3o:DE) yields the primary development-disjoint benchmark of 359 Fabs and 17,816 CDR residues.

We additionally use two external cohorts. A recent panel contains 100 paired antibodies (5,065 CDR residues) released in SAbDab between January 2024 and February 2026 and has no exact-PDB, 90%-identity antibody-family, or 90%-identity antigen-family overlap with the historical collection. A second reserve contains 75 paired antibodies and 3,848 CDR residues whose heavy-chain, antigen, and joint antibody–antigen families are disjoint from development. Together, the three evaluation cohorts contain 534 antibodies and 26,729 CDR residues. Antigen coordinates are not supplied to either expert.

### 4.2 IAF training and comparison methods

All IAF parameters are selected exclusively from the 62-antibody development collection. The static fusion anchor *g*_0_ = 0.59 is obtained by a one-dimensional development-set search over *g* ∈ {0, 0.01, …, 1.00} using mean per-antibody CDR recovery. It serves only as the initialization and regularization anchor for IAF and as a static-fusion baseline.

IAF hyperparameters are selected by five-fold antibody-grouped cross-validation. Within each fold, feature-standardization statistics and the empirical CDR prior are estimated exclusively from the training antibodies. For development targets, empirical-prior features are constructed in a leave-one-target-out manner. The regularization coefficient is selected from {0, 0.01, 0.1, 1} using targetmacro validation NLL, yielding λ = 0.01. After selection, three 4→8→1 IAF models are trained on the complete development collection with seeds 41–43 and frozen; their probability distributions are averaged at evaluation time. Neither pretrained expert is fine-tuned.

We compare IAF with the individual structure and language experts and with static or learned fusion alternatives, including equal-weight and development-calibrated product-of-experts fusion, linear probability mixing, CDR-specific weighting, lower-entropy expert selection, amino-acid residual correction, entropy-based gates, and learned logit-fusion controls. To evaluate dependence on the structural expert, the same IAF training protocol is separately instantiated with AntiFold and AbMPNN. Each refined model is compared with its corresponding frozen structure-only baseline, and no evaluation antibody is used for fitting or recalibration.

### 4.3 Framework interventions and mechanistic analyses

We isolate the contribution of framework identity while keeping the target backbone, structural-expert probabilities, CDR-mask positions, pretrained AntiBERTy checkpoint, and trained IAF models fixed. For each eligible target chain, ten *shuffled-framework* controls replace the target framework with a donor matched by chain role, sequence length, and exact CDR-mask positions. The resulting strictly comparable subset contains 348 development-disjoint antibodies and 16,013 CDR residues. A *no-framework* control masks the complete sequence.

The generic empirical prior *P*_emp_ (*a* | *r, u*) is estimated separately for the six CDR types *r* using ten normalized loop-position bins *u* and a pseudocount of 0.5 per amino acid. During development, the current antibody is excluded when constructing its prior; for evaluation, the prior is constructed once from all 62 development antibodies and frozen.

To localize framework context, we additionally retain either framework residues within eight sequence positions of the evaluated CDR (*local-only*) or the complementary framework positions (*distant-only*). AntiBERTy processes heavy and light chains independently, so all framework-conditioning experiments use same-chain context only.

Mechanistic analyses characterize each residue by structural uncertainty *H* (*P*_*S*_), framework uncertainty *H* (*P*_*F*_), cross-expert disagreement

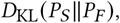

and the informativeness proxy

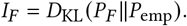

The latter measures departure from a generic development-derived CDR distribution and is not interpreted as a direct estimate of causal framework information; target specificity is established separately by the matched-versus-shuffled intervention. All stratification thresholds are determined from development data and frozen before evaluation.

### 4.4 Evaluation protocol

The primary metric is CDR sequence recovery, calculated within each antibody and then macro-averaged across targets. We additionally report target-macro native negative log-likelihood (NLL) and perplexity. Paired 95% confidence intervals are estimated using 10,000 target-level bootstrap replicates [8]; pooled analyses resample complete targets within each cohort.

We distinguish two decoding settings. In *open-loop* evaluation, IAF refines structural-expert distributions along that expert’s original generated trajectory without altering subsequent structural contexts. In *closed-loop* decoding, the refined argmax residue is written back before the next structural prediction. The framework-conditioned AntiBERTy distributions remain cached and unchanged in both settings. Open loop therefore isolates distribution-level refinement at matched structural contexts, whereas closed loop evaluates the complete autoregressive design process.

We additionally evaluate structural compatibility on the recent 100-antibody panel. Complete paired sequences generated by the base and refined models are folded using the frozen ABodyBuilder2 model-1 checkpoint [1]. Predicted and experimental CDR backbone atoms (N, C*α*, C, and O) are aligned locally by the Kabsch algorithm, and CDR RMSD is reported with paired target-bootstrap confidence intervals. No native CDR identity is supplied during sequence generation or fold-back prediction.

Additional robustness analyses include CDR-specific performance, framework-homology strata, release year and CDR-length groups, residue-level correction transitions, predictive-entropy changes, and sensitivity to fusion settings. Sampling-based comparisons follow the corresponding released model configurations and are reported separately from deterministic open- and closed-loop decoding.

## 5. Results

### 5.1 IAF improves CDR design across structural experts

We first evaluate whether residue-specific information-aware fusion improves two frozen antibody structure experts on the recent 100-antibody panel (Fig. 1). Under the corresponding 20-sample design protocol at *T* = 0.2, AbMPNN reaches 56.50% target-macro recovery, which increases to 64.32% with static fusion and to 65.85% with IAF. IAF therefore improves the frozen AbMPNN baseline by +9.35 percentage points (95% CI +7.43 to +11.48) and, importantly, provides an additional 1.53 points over static fusion (CI +0.93 to +2.14). Native NLL is reduced from 1.4814 to 1.1989.

**Figure 1:**
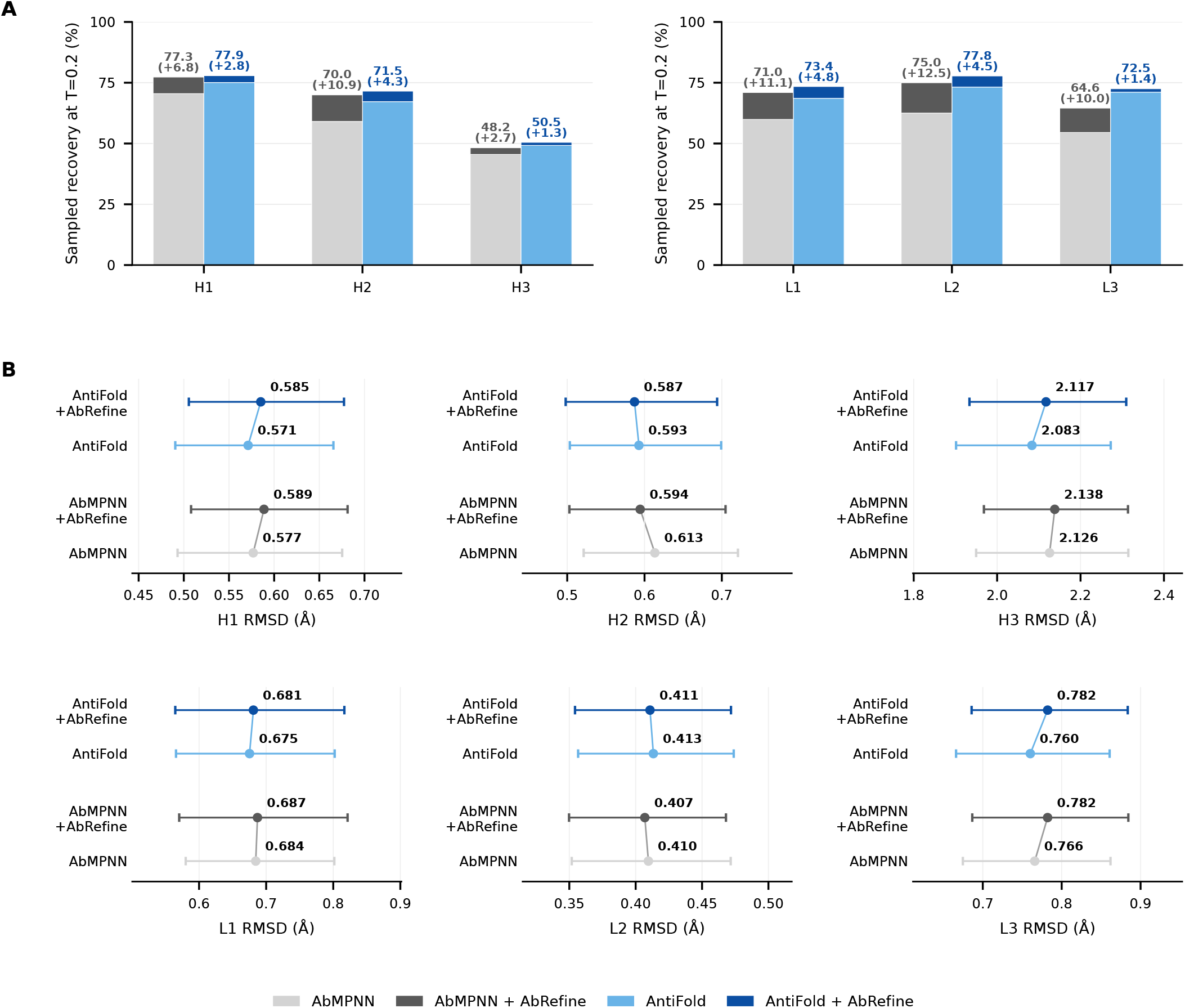
Cross-expert recovery and structural compatibility. (A) Target-macro recovery on the recent 100-antibody panel under the 20-sample, *T* = 0.2 protocol. IAF improves both frozen structural experts, with a significant additional gain over static fusion for AbMPNN. (B) ABodyBuilder2 fold-back evaluation of deterministic greedy designs. Error bars denote paired 95% target-bootstrap confidence intervals.

The stronger AntiFold expert shows a different pattern. Recovery increases from 64.53% to 67.48% with IAF ( +2.95 points; 95% CI +2.07 to +3.93). Relative to static fusion, the additional recovery change is only 0.18 points (CI +0.22 to −0.56), whereas native NLL decreases significantly from 1.1723 to 1.1542. Thus, adaptive fusion provides a clear top-1 advantage when the structural expert leaves greater residual ambiguity, while for the stronger expert its additional benefit is expressed primarily through improved probability assignment.

This contrast is also visible across CDRs. For AbMPNN, IAF exceeds static fusion in five of six regions, with the largest additional gain in H3 ( +2.00 points). For AntiFold, regional changes partially offset one another: H3 improves by +1.13 points, whereas L2 decreases by 1.67 points. The resulting aggregate recovery remains essentially unchanged relative to static fusion despite improved likelihood.

Fold-back evaluation asks whether these sequence-level improvements are accompanied by detectable structural drift (Fig. 1B). Across the six CDRs, the IAF-refined greedy designs show no significant change in aggregate CDR-backbone RMSD relative to either frozen base expert; both paired confidence intervals include zero. The recovery improvements therefore do not produce a detectable loss of compatibility with the prescribed CDR geometry under this computational fold-back assay. This endpoint measures predicted structural consistency rather than binding or experimental function.

### 5.2 IAF generalizes across cohorts and autoregressive feedback

+ + +

We next evaluate the frozen AntiFold-based IAF model on the development-disjoint public benchmark and two external cohorts. On the primary 359-Fab public benchmark, AntiFold reaches 68.91% recovery and IAF reaches 71.17%, a +2.25-point improvement. Relative to static fusion, IAF adds 0.84 points (95% CI +0.51 to +1.18) and reduces native NLL from 1.0599 to 1.0352. All six CDRs improve over the static rule, with the largest additional gain in H3 ( 1.80 points; Table 2).

**Table 1:** IAF performance on the recent 100-antibody panel under the 20-sample, *T* = 0.2 recovery protocol. Recovery is target-macro percent; NLL is evaluated at *T* = 1. Static fusion is included as an ablation of residue-specific adaptation.

| Base expert | Recovery (%) |  |  |  |  | Native NLL |  |  |
| --- | --- | --- | --- | --- | --- | --- | --- | --- |
| | Original | Static | IAF | $\Delta$ vs base | $\Delta$ vs static | Original | Static | IAF |
| AbMPNN | 56.50 | 64.32 | **65.85** | +9.35 | +1.53 | 1.4814 | 1.3161 | **1.1989** |
| AntiFold | 64.53 | 67.30 | **67.48** | +2.95 | +0.18 | 1.3217 | 1.1723 | **1.1542** |

**Table 2:** Open-loop target-macro recovery (%) on the 359-Fab development-disjoint public benchmark.

| CDR | AntiFold | Static | IAF | $\Delta$ vs AF | $\Delta$ vs static |
| --- | --- | --- | --- | --- | --- |
| H1 | 77.14 | 79.23 | 79.82 | +2.68 | +0.59 |
| H2 | 71.56 | 75.61 | 75.84 | +4.28 | +0.23 |
| H3 | 54.39 | 55.16 | 56.96 | +2.57 | +1.80 |
| L1 | 73.40 | 73.54 | 74.33 | +0.93 | +0.79 |
| L2 | 84.72 | 85.31 | 85.77 | +1.06 | +0.46 |
| L3 | 73.94 | 74.25 | 75.13 | +1.19 | +0.88 |
| All | 68.91 | 70.33 | **71.17** | +2.25 | +0.84 |

On the recent external panel and the family-disjoint reserve, IAF reaches 67.93% and 67.29% recovery, corresponding to improvements of +3.08 and +1.78 points over AntiFold. Additional recovery over static fusion is small on these two cohorts ( −0.11 and +0.31 points, respectively), but native NLL is reduced on both. Across the three cohorts, IAF therefore consistently improves the frozen structural baseline, while its advantage over static fusion depends on how much residual ambiguity remains in the underlying expert.

The learned fusion rule also remains stable when its predictions alter subsequent structural contexts. Across the pooled 534-antibody evaluation, open-loop IAF reaches 70.02% recovery and closed-loop decoding reaches 70.10%. The difference is +0.09 points (95% stratified paired-bootstrap CI −0.19 to +0.36), and all six CDR-specific intervals include zero (Fig. 2). Thus, the benefit of residue-specific arbitration is preserved when refined predictions are fed back into the autoregressive structural decoder.

**Figure 2:**
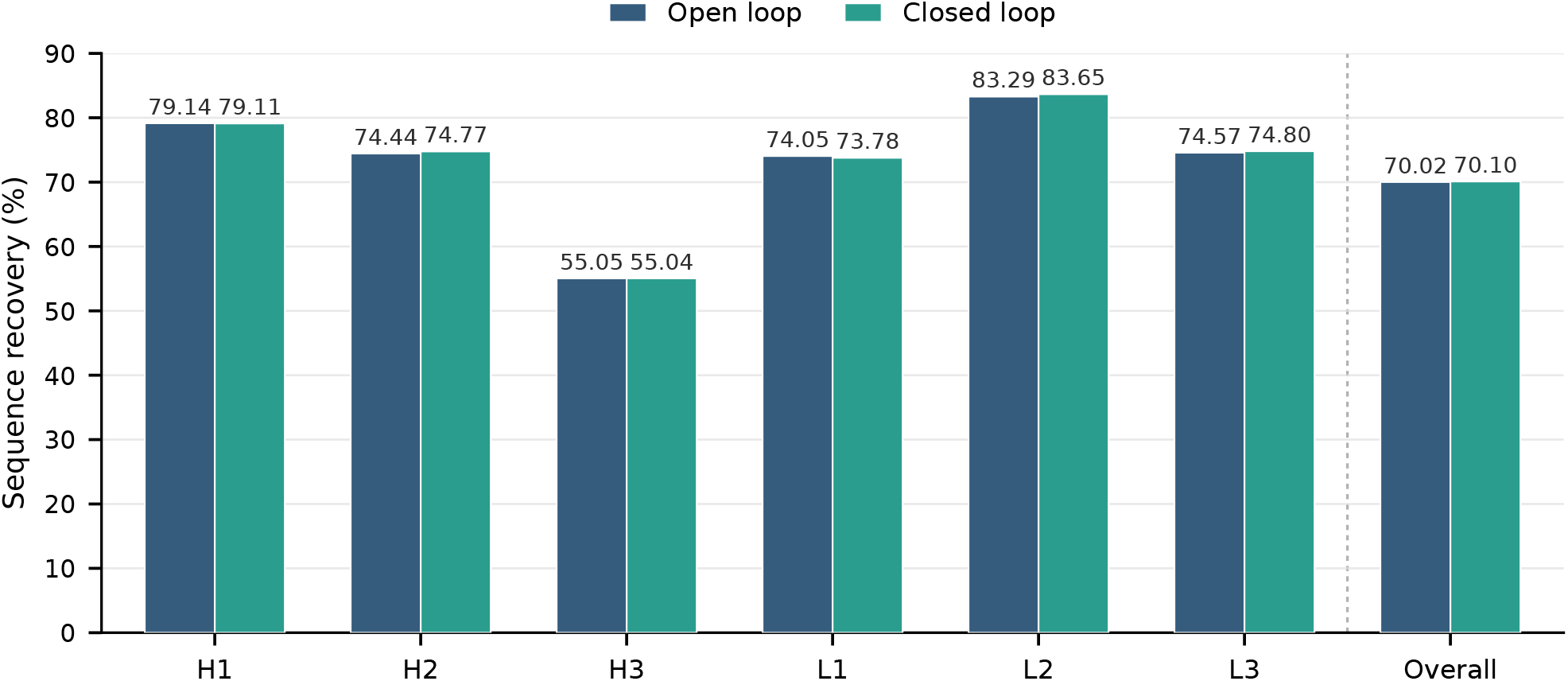
IAF under autoregressive feedback. Open- and closed-loop recovery across the pooled 534-antibody evaluation. Closed-loop feedback changes overall recovery by only +0.09 points (95% stratified paired-bootstrap CI +0.19 to −0.36), with no significant CDR-specific degradation.

### 5.3 The useful auxiliary signal is target-framework specific

Having established predictive gains, we next ask whether the auxiliary sequence signal actually originates from the target framework. We intervene only on the AntiBERTy context while keeping the target backbone, structural predictions, CDR masks, empirical prior, and trained IAF models fixed (Fig. 3A).

**Figure 3:**
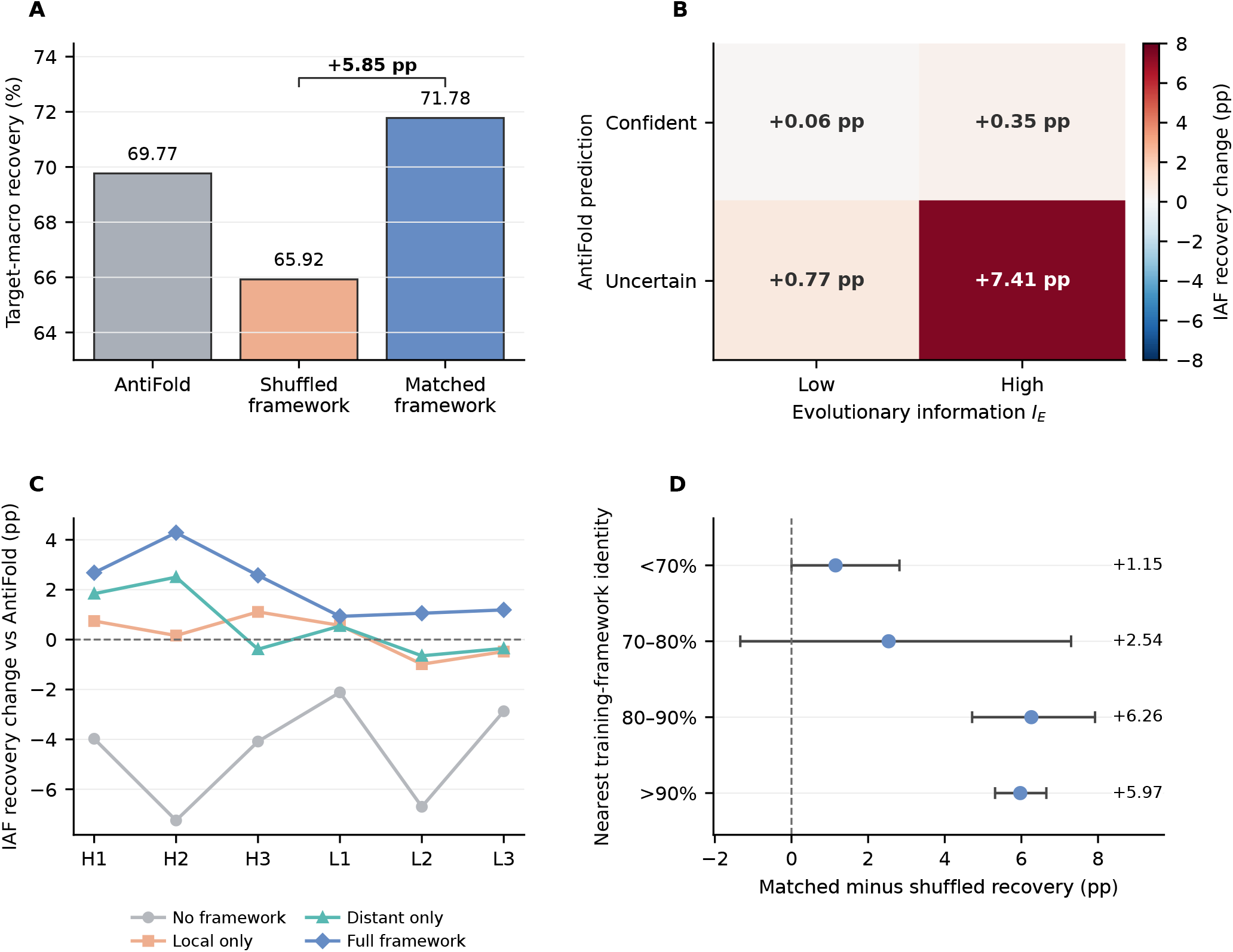
Source and conditional utility of framework information. (A) Target-matched frameworks outperform exact-mask-matched shuffled frameworks while all other model components remain frozen. (B) IAF provides its largest gains where structural uncertainty and framework-distribution informativeness are both high. (C) Neither local nor distant framework context alone reproduces the full-context effect. (D) Matched-over-shuffled effects remain positive across framework-homology strata. Error bars denote paired 95% target-bootstrap confidence intervals.

**Figure 4:**
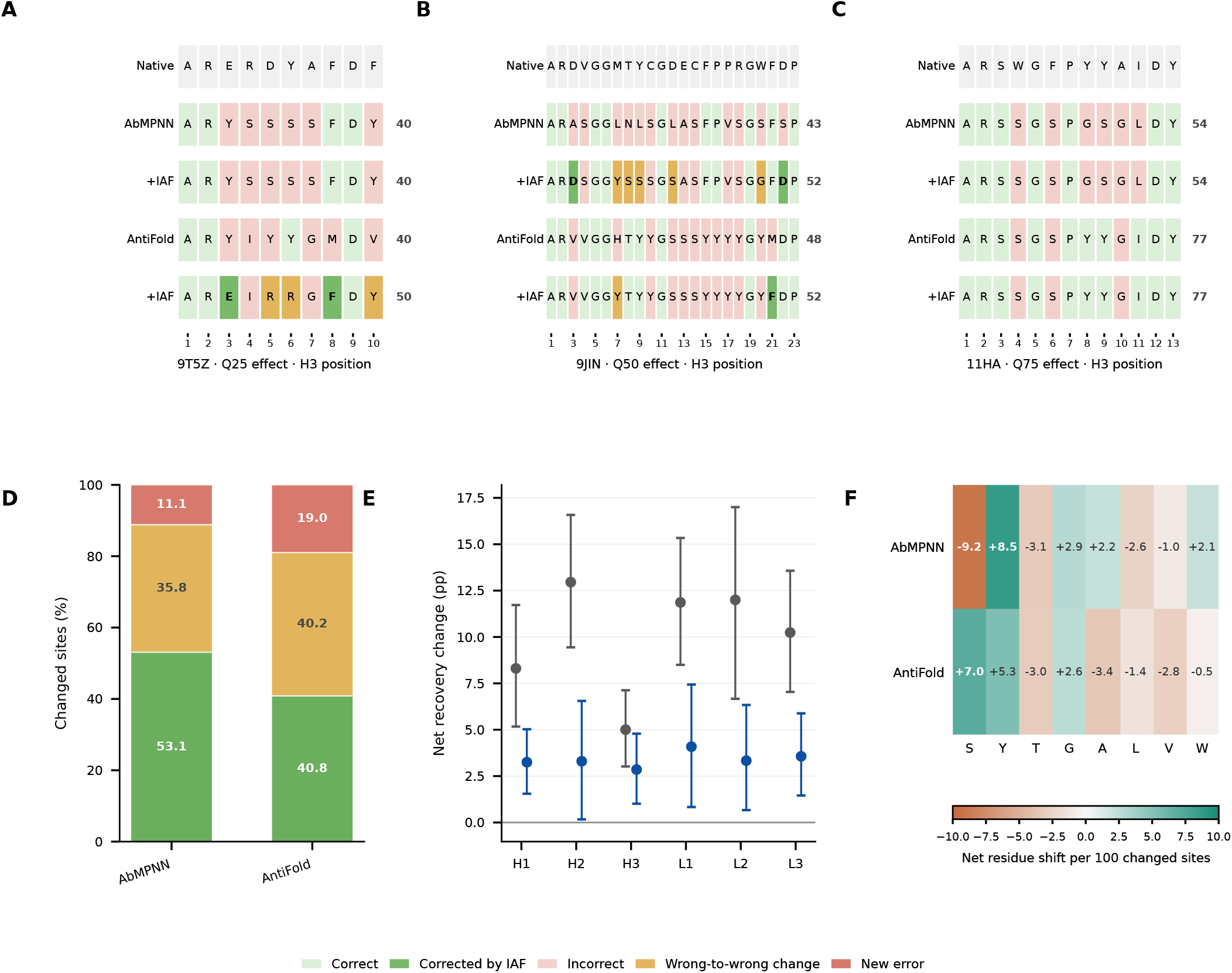
Residue-level effects of IAF. IAF changes a selective subset of structural-model predictions. Corrections substantially outnumber newly introduced errors for both base experts, while amino-acid shifts differ between AbMPNN and AntiFold, indicating base-dependent rather than global compositional correction.

On the 348-target shuffle-compatible subset, AntiFold reaches 69.77% recovery and matched-framework IAF reaches 71.78%. Replacing the target framework with ten donor frameworks matched by chain role, sequence length, and exact CDR-mask positions reduces recovery to 65.92%. Matched context therefore exceeds shuffled context by +5.85 points (95% CI +5.26 to +6.47), with 288 target-level wins, 44 ties, and 16 losses. Importantly, the mean learned framework weight changes only from 0.447 under matched context to 0.441 under shuffling. The performance loss therefore arises primarily from changing the content of the auxiliary distribution rather than from a collapse of the learned gate.

Two additional controls rule out a generic residue-frequency explanation. Removing framework sequence information entirely reduces fused recovery to 58.85%, while a development-derived empirical CDR prior reaches only 68.31%, below the 68.91% AntiFold baseline. The benefit therefore cannot be reproduced by a generic antibody or CDR composition prior.

Framework information is also distributed beyond immediate CDR flanks. Full same-chain context exceeds local-only and distantonly conditioning by + .90 and +1.68 points, respectively (Fig. 3C). The matched-over-shuffled effect remains positive across framework-homology strata, although uncertainty is larger in the sparsely populated low-identity groups (Fig. 3D).

### 5.4 IAF concentrates its benefit where structural evidence is ambiguous

We next examine whether the learned fusion behavior follows the information structure that motivated IAF. Residues are characterized by structural uncertainty and by

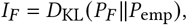

which measures how strongly the framework-conditioned distribution departs from a generic development-derived CDR prior.

When both structural entropy and *I*_*F*_ are high, IAF improves recovery by + 7.41 points (95% CI + 5.53 to + 9.30), with 589 corrected predictions and 231 newly introduced errors (Fig. 3B). In contrast, the other three uncertainty–informativeness quadrants change by only + 0.06, + 0.35, and + 0.77 points, with confidence intervals crossing zero. The gain is therefore highly localized: framework-conditioned evidence is most useful when the structural expert is uncertain and the auxiliary distribution contains information beyond generic CDR preferences.

Residue-level transitions support the same interpretation. On the recent panel, IAF changes 1,113 AbMPNN top-1 predictions, comprising 591 corrections, 124 new errors, and 398 substitutions between non-native residues. For AntiFold, 310 changes are corrections, 144 are new errors, and 305 replace one non-native prediction with another. Corrections therefore outnumber newly introduced errors by 4.77:1 for AbMPNN and 2.15:1 for AntiFold. H3 contains the largest absolute number of changed predictions for both structural experts.

These changes are not explained by a fixed amino-acid composition shift. Among changed sites, AbMPNN loses serine whereas AntiFold gains it, while both models show increases in tyrosine and glycine. IAF therefore responds to the interaction between the structural and framework-conditioned distributions rather than imposing a common residue-frequency correction.

### 5.5 IAF adapts distributional diversity to the structural expert

Finally, we examine how IAF changes the complete predictive distribution rather than only its top-ranked residue. On the recent panel, mean predictive entropy changes from 1.952 to 1.888 bits for AbMPNN (Δ = −0.064; 95% CI −0.135 to +0.003), but from 1.182 to 1.497 bits for AntiFold (Δ = +0.316; CI +0.275 to +0.355; Fig. 5). Thus, the same IAF mechanism slightly concentrates the weaker AbMPNN distribution overall while broadening the stronger AntiFold distribution.

**Figure 5:**
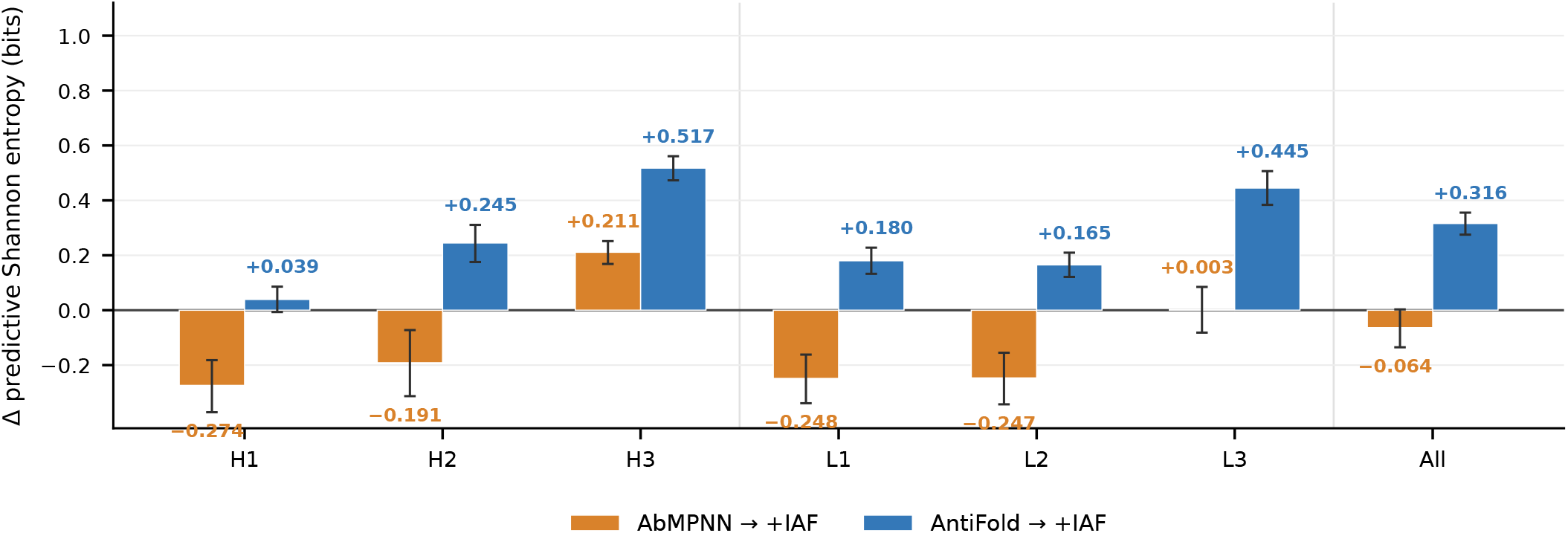
IAF adaptively changes predictive diversity. Refined-minus-base Shannon entropy on the recent 100-antibody panel. Positive values indicate broader amino-acid distributions and negative values indicate concentration. The opposite aggregate effects for AbMPNN and AntiFold show that IAF does not impose a fixed sharpening or smoothing behavior.

The effect is also CDR dependent. For AbMPNN, IAF broadens H3 by +0.211 bits while concentrating H1, H2, L1, and L2. For AntiFold, the largest increases occur in H3 ( +0.517 bits) and L3 ( +0.445 bits), whereas H1 changes by only 0.039 bits. IAF therefore does not implement a universal sharpening operation. Instead, it redistributes probability mass according to the local relationship between structural evidence and framework-conditioned sequence information.

## 6 Conclusion

We introduced AbRefine, an information-aware fusion framework for structure-guided antibody CDR design. Our results show that a prescribed backbone does not exhaust the information available for residue selection: the target framework provides complementary sequence information that helps resolve ambiguity left by structural conditioning. Controlled framework interventions establish that this signal is target specific rather than a generic CDR frequency effect, and its utility is concentrated at positions where structural predictions are uncertain and the framework-conditioned distribution is informative.

AbRefine operationalizes this complementarity through Information-Aware Fusion (IAF), which learns residue-specific corrections to a globally calibrated fusion rule from the information state of two frozen pretrained experts. Across structural experts and evaluation cohorts, IAF improves sequence recovery and native likelihood, remains stable under autoregressive feedback, and provides its clearest advantage over static fusion when the structural expert leaves greater residual ambiguity. Residue-level and entropy analyses further show that IAF does not impose a uniform confidence or amino-acid correction; instead, it adapts the contribution of framework evidence to the local relationship between the two predictive distributions. Computational fold-back evaluation detects no aggregate loss of CDR-backbone compatibility after refinement.

The present study evaluates sequence recovery, likelihood, and predicted structural consistency rather than experimental antibody function. Affinity, specificity, developability, and structural preservation after experimental realization therefore remain to be established, and the current language-model expert is limited to same-chain framework context. Nevertheless, the results support a broader principle for conditional biomolecular design: *geometry constrains the plausible sequence space, contextual sequence information helps resolve the remaining ambiguity, and information-aware fusion determines how strongly that information should be used*.

## A. Additional Protocol Details

### A.1 Input masking and decoding contract

Native CDR identities are excluded from every model input. For each antibody chain, all annotated CDR residues are masked simultaneously before AntiBERTy inference: H1, H2, and H3 are masked together on the heavy chain, and L1, L2, and L3 on the light chain. The resulting framework-conditioned distributions are computed once per chain and cached throughout decoding. AntiBERTy receives neither backbone coordinates nor generated CDR residues.

The structural expert begins with all design positions hidden and receives only the prescribed backbone, observed framework, and the generated prefix permitted by the decoding protocol. Native CDR residues are joined to predictions only after inference for retrospective evaluation.

Each per-residue prediction record stores target and CDR identifiers, the complete 20-way predictive distribution, the native residue used for scoring, and an explicit native_CDR_tokens_visible=false assertion. The evaluation pipeline rejects duplicate residue identifiers, inconsistent annotations, unmatched method pairs, and any record that violates this assertion.

In open-loop evaluation, IAF refines structural-expert distributions recorded along the base expert’s original decoding trajectory; the refined prediction does not modify subsequent structural contexts. In closed-loop decoding, the IAF argmax residue is written into the generated prefix before the next structural prediction. The cached framework-conditioned distribution remains unchanged in both protocols.

### A.2 Development-only IAF training

All quantities used by IAF are determined exclusively from the 62-antibody development collection. The static fusion anchor *g*_0_ = 0.59 is selected by a one-dimensional search over

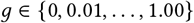

using mean per-antibody CDR recovery. This coefficient serves as the initialization and regularization anchor for IAF and as a static-fusion comparison; it is not the final adaptive prediction rule.

IAF hyperparameters are selected by five-fold antibody-grouped cross-validation, with complete antibodies retained within folds. For each fold, the means and standard deviations used to standardize the four IAF features are estimated exclusively from the training antibodies. The regularization coefficient is selected from

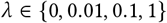

using target-macro validation NLL, yielding λ = 0.01.

The empirical CDR prior is estimated separately for H1, H2, H3, L1, L2, and L3 using ten normalized loop-position bins and a pseudocount of 0.5 for each amino acid. During development, the prior used for a target is constructed after excluding that complete antibody. For evaluation targets, the prior is reconstructed once from all 62 development antibodies and then frozen. Consequently, neither the empirical prior nor the derived informativeness feature contains sequence information from an evaluation antibody.

After model selection, three independent 4→8→1 IAF networks are trained on the complete development collection with seeds 41–43. Each network is optimized independently with the target-macro NLL and anchor regularization defined in the main text. Their probability outputs, rather than their gate values, are averaged at inference time. The structural and language experts remain frozen throughout training and evaluation.

The development-derived feature-normalization statistics, empirical prior, *g*_0_, λ, network architecture, and three trained IAF parameter sets are frozen before any reported evaluation.

### A.3 Target-level metrics and paired inference

For target *t* with evaluated residue set *I*_*t*_, target recovery and native negative log-likelihood are

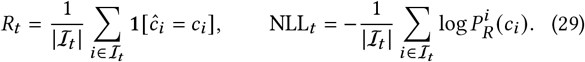

Reported recovery and NLL are macro-averaged across complete antibody targets,

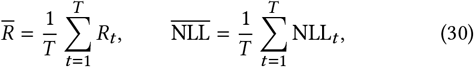

and perplexity is exp 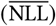 . CDR-specific results use the same procedure after restricting _*t*_ to the corresponding region.

All confidence intervals are paired at the antibody level. We generate 10,000 bootstrap replicates by resampling complete targets with replacement and recomputing the paired method difference. Residues are never treated as independent bootstrap units. For pooled analyses, targets are resampled within each evaluation cohort and the resampled cohorts are recombined at their observed sizes.

### A.4 Public-benchmark provenance and coverage

The public split files correspond to Zenodo 8164693. After deterministic removal of duplicate public rows, the released SAbDab partition contains 2,922 training, 390 validation, and 405 test Fab units, with no 90%-identity six-CDR cluster shared across partitions. Local reconstruction recovers usable paired heavy–light records for 362 test units; 43 units lack at least one required chain record.

For direct compatibility with the reconstructable public split, we retain the 362-target result as a secondary reference. The primary public evaluation additionally excludes 5w3l:EG, 5w3m:EG, and 5w3o:DE because their PDB identifiers occur in the 62-antibody development collection. The resulting exact-PDB-disjoint benchmark contains 359 paired Fabs and 17,816 CDR residues.

Because the released unit is a Fab rather than a PDB entry, distinct Fabs from the same PDB can occur in different sequence-defined partitions. The AntiFold checkpoint SHA256 begins d5c442fa, and the AntiBERTy checkpoint SHA256 begins f1ae33ea. Complete checkpoint digests, target lists, exclusions, and per-residue prediction records accompany the reproducibility package.

## B. Additional Results

### B.1 Family-disjoint reserve

The family-disjoint reserve provides an evaluation in which antibody and antigen families are separated from development. IAF improves both open- and closed-loop recovery over AntiFold and reduces native NLL (Table 3). The additional recovery relative to static fusion is modest, consistent with the observation in the main text that adaptive weighting provides its largest top-1 advantage when the structural expert leaves greater residual ambiguity.

**Table 3:** Target-macro performance on the 75-antibody family-disjoint reserve. Static denotes the development-calibrated distribution-fusion baseline used to anchor IAF.

| Method | Recovery (%) | Native NLL |
| --- | --- | --- |
| AntiFold | 65.51 | 1.3108 |
| Static, open loop | 66.98 | 1.1649 |
| Static, closed loop | 66.51 | 1.1925 |
| IAF, open loop | **67.29** | **1.1480** |
| IAF, closed loop | 67.09 | 1.1614 |

### B.2 Cross-cohort open- and closed-loop IAF evaluation

Table 4 reports IAF separately for the three evaluation cohorts. Open- and closed-loop recovery are similar in every cohort, and none of the paired closed-minus-open confidence intervals excludes zero. This supports the main-text conclusion that residue-specific fusion remains stable when refined predictions alter subsequent structural contexts.

**Table 4:** Frozen-IAF open- and closed-loop results. Confidence intervals compare closed-with open-loop recovery using paired target bootstrap; the pooled interval uses cohort-stratified resampling.

| Cohort | Targets | Open rec. | Closed rec. | Closed-open [95% CI] | Open NLL | Closed NLL |
| --- | --- | --- | --- | --- | --- | --- |
| Public split | 359 | 71.17 | 71.27 | +0.11 [−0.21, +0.43] | 1.0352 | 1.0402 |
| Recent external | 100 | 67.93 | 68.15 | +0.23 [−0.43, +0.89] | 1.1542 | 1.1421 |
| Family reserve | 75 | 67.29 | 67.09 | −0.20 [−1.03, +0.70] | 1.1480 | 1.1614 |
| Pooled | 534 | 70.02 | 70.10 | +0.09 [−0.19, +0.36] | 1.0734 | 1.0763 |

### B.3 Entropy-only gating does not explain IAF To test whether structural uncertainty alone is sufficient for adaptive fusion, we compare global, hard-threshold, and differentiable entropy-gated fusion under identical target-grouped folds. This analysis is diagnostic only and is not used to select any reported evaluation model

Neither entropy-gated variant improves recovery over global fusion. Together with the main-text uncertainty–informativeness analysis, this indicates that uncertainty identifies where refinement may be useful but is insufficient by itself to determine whether the framework-conditioned expert provides corrective information. IAF therefore conditions on the joint information state rather than structural entropy alone.

### B.4 Static-fusion sensitivity

The static fusion rule used to initialize and regularize IAF is not sensitive to a narrowly tuned coefficient. Development recovery is 73.48%, 73.59%, and 73.47% at *g* = 0.58, 0.59, and 0.60, respectively, and a broad range of coefficients remains above the AntiFold baseline on the recent panel. This analysis motivates treating *g*_0_ = 0.59 as a stable global anchor rather than a biologically meaningful constant.

For completeness, Fig. 6 summarizes the behavior of the static baseline before residue-specific adaptation. Recovery gains increase with structural uncertainty, while the broad weight-sensitivity curve motivates learning local deviations around a stable global operating point.

**Figure 6:**
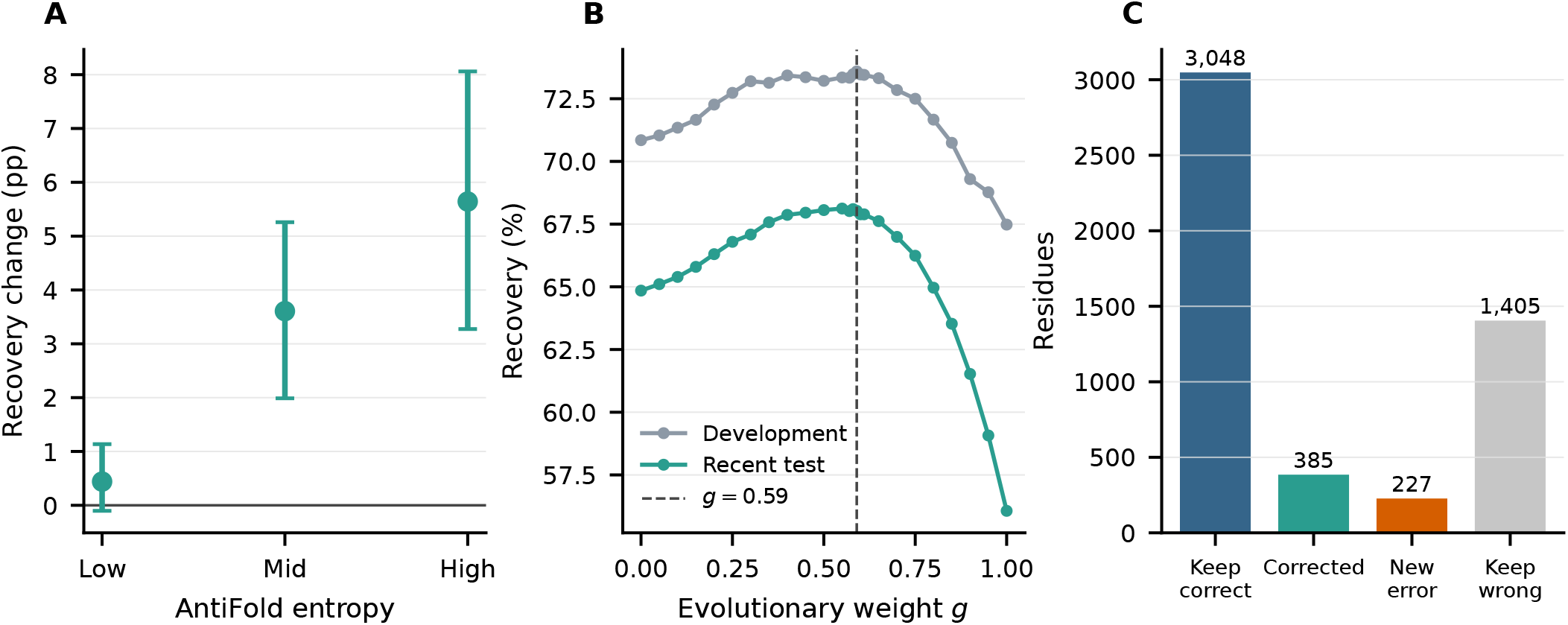
Static-fusion diagnostics. (A) Recovery gains increase across AntiFold entropy strata. (B) Development and descriptive recent-panel sensitivity to the global fusion coefficient; the dashed line marks the IAF anchor *g*_0_ = 0.59. (C) Residue-level transitions under closed-loop static fusion. These analyses characterize the global baseline from which IAF learns residue-specific corrections.

**Figure 7:**
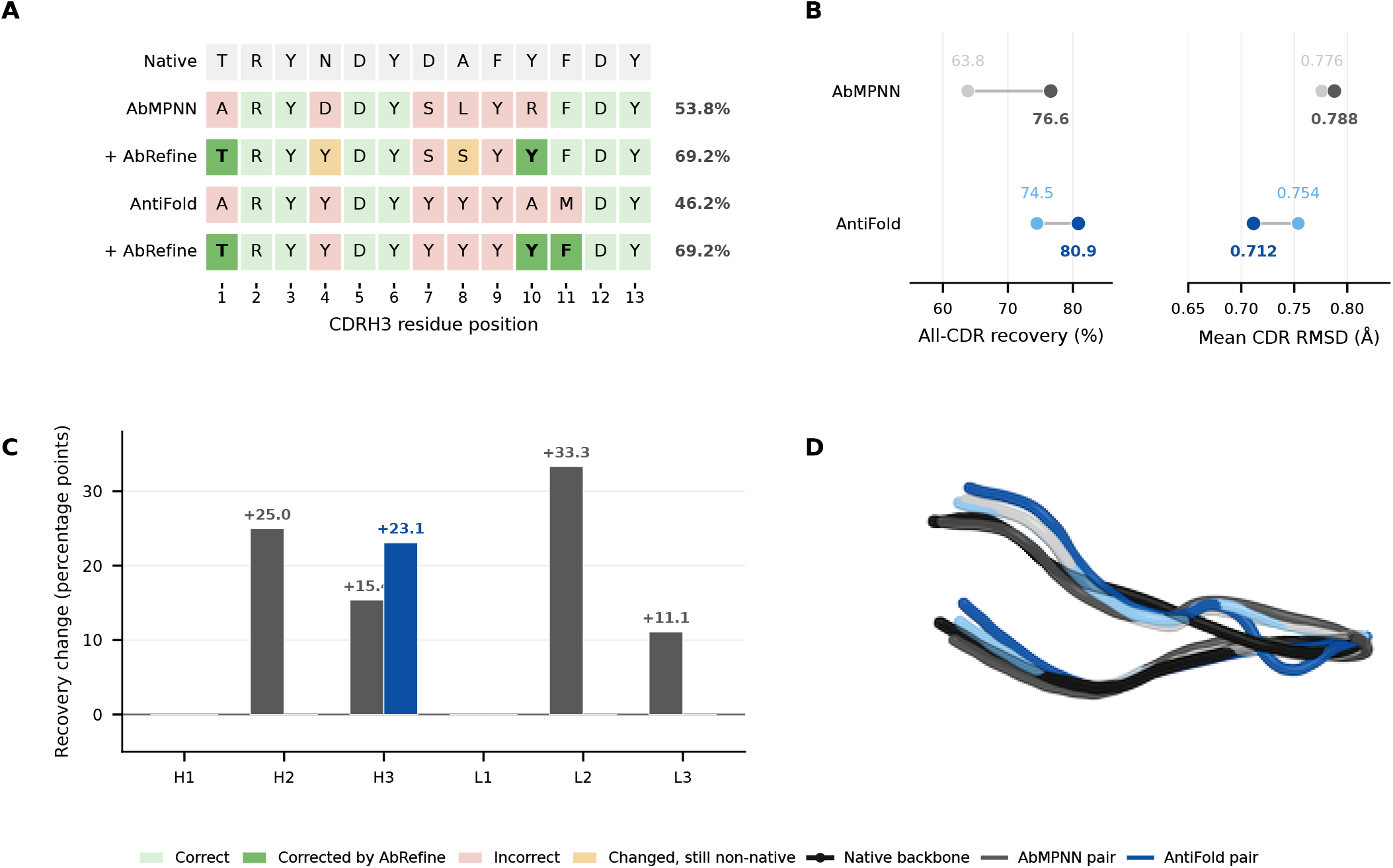
Representative fixed-backbone CDR design for PDB 9PY6. The target is selected by a prespecified median-effect rule among structurally stable cross-base responders. Native residues are used only for retrospective evaluation. The structural panel shows locally superposed experimental and predicted CDRH3 backbones after N/C*α*/C/O alignment.

### B.5 Representative application case

We select a representative application case using a prespecified rule rather than choosing the largest observed improvement. Among recent-panel antibodies with positive recovery gains for both structural experts and at most 0.05 Å absolute mean-RMSD change for either base, PDB 9PY6 lies at the median summed cross-base recovery gain.

With all 47 native CDR residues hidden, IAF raises recovery from 63.83% to 76.60% for AbMPNN and from 74.47% to 80.85% for AntiFold. Mean six-CDR fold-back RMSD changes from 0.776 to 0.788 Å for AbMPNN and from 0.754 to 0.712 Å for AntiFold. The example illustrates how the same framework-conditioned fusion mechanism can correct different residue errors produced by distinct structural experts without selecting an extreme responder.

